# Intraperitoneal treatment with antimicrobial peptide rescues mice from a pulmonary *Francisella* infection

**DOI:** 10.1101/603233

**Authors:** Monique L. van Hoek, Akanksha Kaushal, Barney M. Bishop, Stephanie M. Barksdale

## Abstract

Our long-term goal is to identify new antimicrobial peptides that might be effective against pneumonic *Francisella* infection in mice. Previously, our group searched the peptidome of the American alligator for novel cationic antimicrobial peptides and identified a naturally-occurring C-terminal fragment of apolipoprotein C-1, which we called Apo6. This peptide was found to have antibacterial activity against the ESKAPE pathogens, including those exhibiting multi-drug resistance. In this work, we tested Apo6 and synthetic derivatives for antibacterial activity against *Francisella tularensis* including the virulent strain *F. tularensis* SchuS4. *Francisella* is inherently highly resistant to the cyclic peptide polymyxin antibiotics and beta-lactam antibiotics. We found that our synthetic peptide derivatives (called GATR peptides), designed with increased hydrophobicity and charge, had generally stronger *in vitro* antimicrobial activity against *Francisella* than the parent peptide Apo6. The GATR peptides had a greater effect on the bacterial membrane than the Apo6 peptide and were able to bind *Francisella* LPS, suggesting their mechanism of action against *Francisella*. Cytotoxicity experiments showed low cytotoxicity for most of the GATR peptides, and whole organism toxicity studies in the waxworm allowed us to down-select to two our lead peptides, GATR-3 and GATR-6. These peptides were tested in a murine pulmonary tularemia model. We found that the GATR-3 peptide rescued 50-60% of mice from lethal tularemia infection when administered systemically through the intraperitoneal route. This peptide is a candidate for further pre-clinical studies for a potential peptide-based approach to tularemia.

## Introduction

*Francisella tularensis* is a Gram-negative bacterium that is the causative agent of tularemia. The virulent strains (*F. tularensis tularensis*) can cause disease in humans with inhalation of as few as 10 organisms. In addition, this organism is easily aerosolized and has historically been developed as a bioweapon [1–3]. The United States government has classified *F. tularensis* as a Tier 1 and Category A Select Agent. *F. tularensis* subsp. *tularensis* is found in the United States [4], with localized outbreaks currently occurring across the continent, and is also referred to as the Type A strain. The less virulent Type B strain (*F. tularensis* subsp. *holarctica*) is more commonly found to infect humans in Europe [5]. *F. tularensis* infections (tularemia) are normally treated with fluoroquinolones and aminoglycosides, but are inherently resistant to some antibiotics such as beta-lactams [3, 6] and polymyxins [7, 8]. In addition, drug resistance to conventional antibiotic treatments may be emerging in this species [9, 10], and there is a concern about potentially engineered resistance in the biothreat context. Because of this, there is interest in developing new potential treatments for tularemia [3].

Cationic antimicrobial peptides are small positively-charged peptides, some of which are produced by the innate immune system of vertebrates as well as other organisms [11]. Cryptic cationic antimicrobial peptides are proteolytic fragments of larger proteins; these larger proteins have annotated functions that are not themselves antimicrobial [12, 13]. Cationic antimicrobial peptides can have broad or specific activity against bacteria, viruses, or fungi and can also have host-directed immuno-modulatory functions [11, 14, 15]. Our group has been working to discover novel antimicrobial peptides from reptiles, including the American alligator (*Alligator mississippiensis*) and the Komodo dragon (*Varanus komodoensis*) [16–18]. We have discovered a number of antimicrobial peptides with strong antimicrobial activity against a range of bacteria. Previously, we found that peptides representing the C-terminal fragments of an apolipoprotein in *A. mississippiensis* had antimicrobial activity against *Bacillus cereus*, *Escherichia coli*, *Pseudomonas aeruginosa*, and *Staphylococcus aureus*, as well as drug-resistant strains of *E. coli, P. aeruginosa, S. aureus*, and *Acinetobacter baumannii* [16, 19].

Previous work using canonical and synthetic antimicrobial peptides delivered to the mouse lung as a potential therapy for pneumonic tularemia in mice [20] was not able to demonstrate any significant rescue from infection. A variety of peptides were tested including the human cathelicidin peptide, LL-37, and synthetic fusions of peptides. We sought to identify antimicrobial peptides that might be effective against pneumonic *Francisella* infection in mice.

In this work, we tested one of our previously discovered alligator peptides, called Apo6, against *Francisella*. We then created a set of synthetic peptide derivatives of the Apo6 peptide by making amino acid substitutions that increased the hydrophobicity and charge of the peptides in order to increase the potency. These synthetic peptides are named GATR-1 through GATR-7. We examined how these variations change the mechanism of action and the binding to *Francisella* lipopolysaccharide (LPS). We down-selected the set of peptides based on cytotoxicity, and tested the efficacy of our lead peptides in two infection models: first in the waxworm (*Galleria mellonella*) and then in a murine tularemia pneumonic infection model.

## Materials and Methods

### Bacterial strains

*Francisella tularensis* subsp. *holarctica* CDC Live Vaccine Strain (NR-646), *F. tularensis* subsp. *tularensis* NIH B38 (NR-50), an attenuated strain, and *F. tularensis* subsp. *tularensis* SchuS4 (NR-10492), the fully virulent strain, were obtained from BEI Resources (Manassas, VA). Bacteria were grown 48-72 h on chocolate II agar (BD 211267) at 37°C with 5% CO_2_. Prior to the experiments below, bacteria were scraped off the plate and resuspended to 0.5 McFarland units in phosphate buffered saline (PBS) or Buffer Q [6.12 mM sodium monohydrogen phosphate heptahydrate; 3.92 mM monosodium phosphate anhydrous; 0.3 g/L tryptic soy broth (BD211825); 1 mg/L cysteine HCl]. A standard curve of bacteria was used to determine the CFU equivalents (0.5 McFarland units = 1 × 10^7^ CFU/ml). Resuspended bacteria were then diluted to the appropriate concentration needed. All work with *F. tularensis* SchuS4 was performed in a BSL-3 laboratory following strict safety guidelines.

### Peptide synthesis

Peptides were synthesized by ChinaPeptides, Inc (Shanghai, China) using Fmoc chemistry. Peptide was provided at >95% purity, which was confirmed with RP-HPLC and ESI-MS. Sequences and physico-chemical properties are shown in **Table 1**.

### Peptide properties

Physico-chemical properties including charge, hydrophobic moment and hydrophobicity as well as helical wheels were calculated using Heliquest [21]. In addition, the APD defined total hydrophobic ratio was calculated with the APD3 website [22].

### Minimal inhibitory concentration (MIC) determination assay

MICs were determined according to CLSI guidelines for this organism [23, 24]. Briefly, *Francisella* bacteria were grown on chocolate II agar (BD 221169) for 48-72 h prior to experiments. Minimal inhibitory concentration experiments were performed in Cation-adjusted Mueller Hinton Broth (BD 212322, CAMHB) with 2% IsovitaleX (BD 211875) using polypropylene plates [25]. Approximately 3 × 10^4^ bacteria were added to each well, as determined using a McFarland standard curve for *Francisella*. Results were analyzed at 21 and 42 µg/ml using a two-way ANOVA with Sidak’s multiple comparisons.

### Antimicrobial assays

The antimicrobial activity of antimicrobial peptides against *F. tularensis* was determined as described previously [26–28]. Briefly, in a 96 well plate, 1 × 10^5^ CFU per well were incubated with various peptide concentrations in Buffer Q for 3 h at 37°C (total volume 100 µl). After the incubation, well contents were serially diluted, and 5 µl of each dilution was spotted onto chocolate agar and allowed to dry. Agar plates were incubated overnight at 37 °C and the colonies were counted. The concentration of peptide required to kill 50% of microbial population (EC_50_) was analyzed by analyzing the percentage of surviving colonies after the overnight incubation as a function of log of peptide concentration. The data was analyzed through GraphPad Prism 6 (GraphPad Software Inc. San Diego, CA, USA). The antimicrobial activity of the derivatives was compared to the activity of LL-37, a human cathelicidin with known antibacterial activity against *Francisella* [27, 29]. The confidence intervals along with the EC_50_ values for each peptide are reported in Table 2. Samples were run in triplicate on three separate occasions.

**Table 2.** Antibacterial activity of GATR peptides against *F. tularensis* LVS and NIH B38. EC_50_ values of Apo6 and its derivatives were determined in Buffer Q against *F. tularensis* LVS *and F. tularensis* NIH B38 (the Type strain). For statistical comparison, the 95% confidence intervals (p < 0.05) are listed. The values are also expressed as µM for direct comparison.

|  | *F. tularensis* LVS |  | *F. tularensis* NIH B38 |  |
| --- | --- | --- | --- | --- |
| Peptide name | EC<sub>50</sub> (95% CI) [µg/ml] | EC<sub>50</sub> [µM] | EC<sub>50</sub> (95% CI) [µg/ml] | EC<sub>50</sub> [µM] |
| Apo6 | 6.8 (5.9-7.8) | 2.5 | 16 (8.6-31) | 5.89 |
| GATR-1 | 0.76 (0.54-1.1) | 0.26 | 16 (9.5-27) | 5.62 |
| GATR-2 | 2.4 (1.5- 3.7) | 0.86 | 11 (7.7-15) | 3.9 |
| GATR-3 | 0.53 (0.42-0.66) | 0.19 | 0.80 (0.56-1.1) | 0.28 |
| GATR-4 | 11.0 (7.5-16) | 3.7 | Not active | Not active |
| GATR-5 | 0.76 (0.56- 1.0) | 0.26 | 1.9 (1.6-2.3) | 0.65 |
| GATR-6 | 0.89 (0.60-1.3) | 0.29 | 0.16 (0.06-0.43) | 0.051 |
| GATR-7 | 1.0 (0.81-1.3) | 0.32 | 1.7 (1.4-2.1) | 0.53 |
| LL-37 | 0.21 (0.14-0.31) | 0.047 | 0.13 (0.076-0.21) | 0.028 |

### Membrane depolarization assay

Membrane potential was measured using a fluorescent assay utilizing DiSC_3_(5) dye as previously described with some modification [19, 28]. *F. tularensis* LVS was grown on chocolate II agar (48 h, 37°C, 5% CO_2_), and the colonies were suspended in 10 mM phosphate buffer to 0.5 McFarland standard. 100 µL of this suspension was added to wells of a black polypropylene 96 well plate. The plate was incubated in a Tecan Infinite F200 fluorimeter. A change in the fluorescence was monitored until equilibrium is reached, evidenced by quenching of the fluorescent signal, indicating maximum uptake of the dye by the membrane. The experimental wells were then treated with 100 µl of various concentrations of peptide diluted in 10 mM phosphate buffer. The plate was returned to the spectrofluorometer and readings were taken every min for 15 min (excitation=620 nm; emission=670 nm). Peak RFU at each concentration was used in the analysis. Samples were run in triplicate on two separate occasions. Bacteria without peptide treatment were used as a negative control, and LL-37 was used as positive control. Depolarization results were analyzed using a one-way ANOVA with Dunnett’s multiple comparisons.

### Ethidium bromide uptake assay

Pore formation in *F. tularensis* LVS cytoplasmic membrane was assessed using ethidium bromide as described previously with some modification [19, 28]. *tularensis* LVS was grown on chocolate II agar (48 h, 37°C, 5% CO_2_) and colonies scraped into solution. Bacteria were suspended in 10 mM phosphate buffer to 0.5 McFarland standard. In a black polypropylene 96 well plate, 180 µL bacterial culture was then mixed with 10 µM ethidium bromide (final concentration) and incubated with varying concentrations of peptide. The plate was read in a Tecan infinite F200 fluorimeter every 2 min for 20 min at 37°C (excitation=535 nm, emission=590 nm). Data shown is from the 20 min mark. Samples were run in triplicate on three separate occasions. Bacteria without peptide was used as a negative control, and LL-37 was used as positive control. Results were analyzed using a one-way ANOVA with Dunnett’s multiple comparisons.

### LPS binding

To examine the potential binding between *F. tularensis* LVS lipopolysaccharide (LPS) and the GATR peptides, an LPS-binding assay using 1,9-dimethylmethyl blue (DMMB) was performed as previously described [30]. LPS from *F. tularensis* subsp. *holarctica* LVS was obtained from BEI Resources (NR-2627). Briefly, 150 µg/ml of LPS was incubated with 10 µg/ml of peptide in distilled endotoxin-free water for 1 h. The solution was added to DMMB, and the absorbance was read at 535 nm on a spectrometer. Samples were run in triplicate on two separate occasions. Results were analyzed using a one-way ANOVA with Dunnett’s multiple comparisons.

### Hemolysis assay

The hemolysis assay was performed using washed, defibrinated sheep blood as previously described [31]. Sheep red blood cells (2% RBC) in phosphate buffered saline (PBS) were added to various dilutions of peptide reconstituted in PBS in a sterile U-bottom 96 well plate. The plate was incubated for 1 h at 37 °C and then centrifuged at 1000 rpm for 2 min. The supernatant was transferred to a fresh plate and read at 540 nm on a spectrometer. Sheep RBCs (2%) with PBS alone served as the negative control, and 2% RBC in water as the positive control. Experiment was performed twice in triplicate. A representative experiment is shown Results were analyzed using a one-way ANOVA with Dunnett’s multiple comparisons.

### Cytotoxicity assay

Cytotoxicity assays were performed using the Vybrant MTT Cell Proliferation Assay Kit (Life Technologies) according to manufacturer’s instructions. Assays were performed using human lung epithelial carcinoma line A549 (ATCC CCL-185) and human liver carcinoma line HepG2 (ATCC HB-8065), which were maintained at a low passage in Dulbecco’s Minimal Essential Media (Life Technologies 11995073) with 10% heat-inactivated fetal bovine serum and 13 U/ml penicillin-streptomycin. 100 µg/ml of peptide was used for each experimental well, added to the cell growth medium, and incubated for 24 h. Each experiment was performed in triplicate two times. A representative experiment is shown. Results were analyzed using a one-way ANOVA with Dunnett’s multiple comparisons.

### Peptide toxicity in *Galleria mellonella* larvae

Larvae were used to assess *in vivo* toxicity of peptides. *G. mellonella* larvae (greater wax moth larvae or “waxworms”) were obtained from Vanderhorst Wholesale (Saint Marys, OH, USA). Ten larvae of equal size/weight were randomly assigned to each group and placed into labeled petri dishes. A 1 ml syringe with a 27G needle was used to inject 10 µl containing 10 µg peptide into each larvae’s right proleg. Survival was observed for 48 h. Results from one representative experiment of two total are shown and were analyzed using a Mantel-Cox test.

### *G. mellonella* infection and treatment

Survival assay of wax moth larvae following *Francisella* infection with and without treatment was conducted as previously described [2, 32, 33]. *G. mellonella* (wax moth larvae or waxworms) were obtained from Vanderhorst Wholesale (Saint Marys, OH, USA). Ten larvae of equal size/weight were randomly assigned to each group and placed into labeled petri dishes. A 1 ml syringe with a 27G needle was used to inject 10 µl of 1×10^8^ CFU/ml of *F. tularensis* LVS into each larvae’s right proleg. After a 60 min incubation to allow the infection to occur, the larvae were then injected with 10 ul of either PBS (no treatment) or 10 ng of the derivatives in the larvae’s left proleg. Bacteria treated with 10 µg of levofloxacin was used as a positive control. The experiment was conducted twice; one representative experiment is shown.

### Animal model of tularemia infection

Female BALB/c mice 6-8 weeks of age were obtained from Jackson Laboratories. Animal experiments were approved by and conducted in compliance with regulations of the Institutional Animal Care and Use Committee (Protocol #0328) of George Mason University. All experiments were carried out in accordance with the National Research Council’s Guide for the Care and Use of Laboratory Animals (2011) and the Public Health Service Policy on Humane Care and Use of Laboratory Animals (2002). Animals were scored twice daily based on appearance, activity, respiration, and appearance following our protocol. If mice were weighed, weights were taken individually prior to any experimental work each day.

For the inoculum, *F. tularensis* LVS was grown for 2 days on chocolate II agar (37°C, 5% CO2). Colonies were scraped and resuspended in sterile PBS to 0.5 McFarland Standard (which is equivalent to ∼10^7^ CFU/ml for this organism). 36 µl of this suspension was added to 10 ml of sterile PBS, and dilution plating was subsequently performed to confirm the inoculation dose. Mice were lightly anesthetized using isoflourane immediately before infection. Each mouse received an intranasal inoculation of 25 µl of this secondary suspension, evenly divided between both nares.

Peptide treatments were performed through intraperitoneal (IP) injections. Each injection consisted of 500 µl PBS containing 100 µg peptide or 60 µg levofloxacin. Treatments were performed 3 h, 24 h, and 48 h after infection. In addition, one group (5 mice) received a prophylactic treatment 24 h before infection in the first study, and one group received no treatment. Survival was tracked for 13 days. Survival results were analyzed using a Mantel-Cox test.

For organ burden studies, mice were infected and treated as above and sacrificed on Day 4. Lungs, livers, and spleens were harvested and homogenized in PBS using DT-20 tubes with an ULTRA-TURRAX Tube Drive (IKA, Wilmington, NC, USA). Homogenate was plated on chocolate II agar and incubated for two days (37 °C, 5% CO2). CFU counts were analyzed using a one-way ANOVA with Dunnett’s multiple comparisons.

### Statistical analysis

All statistical analysis was performed in GraphPad Prism 6.0 or 7.0. Tests performed are listed in each methods section and figure legend.

## Results

### Peptide design and properties

Apo6 is a naturally occurring (native) peptide identified intact from American alligator blood by *de novo* peptide mass-spectrometry sequencing [16, 17]. It is the C-terminal sequence of alligator apolipoprotein E, and was discovered using our BioProspector process [16].

A series of Apo6 derivative peptides, designated GATR-1 through GATR-7 (Table 1), were generated by introducing changes in the original Apo6 sequence, in order to improve the peptide’s amphipathicity, hydrophobic face, or net charge, as described below.

**Table 1.**
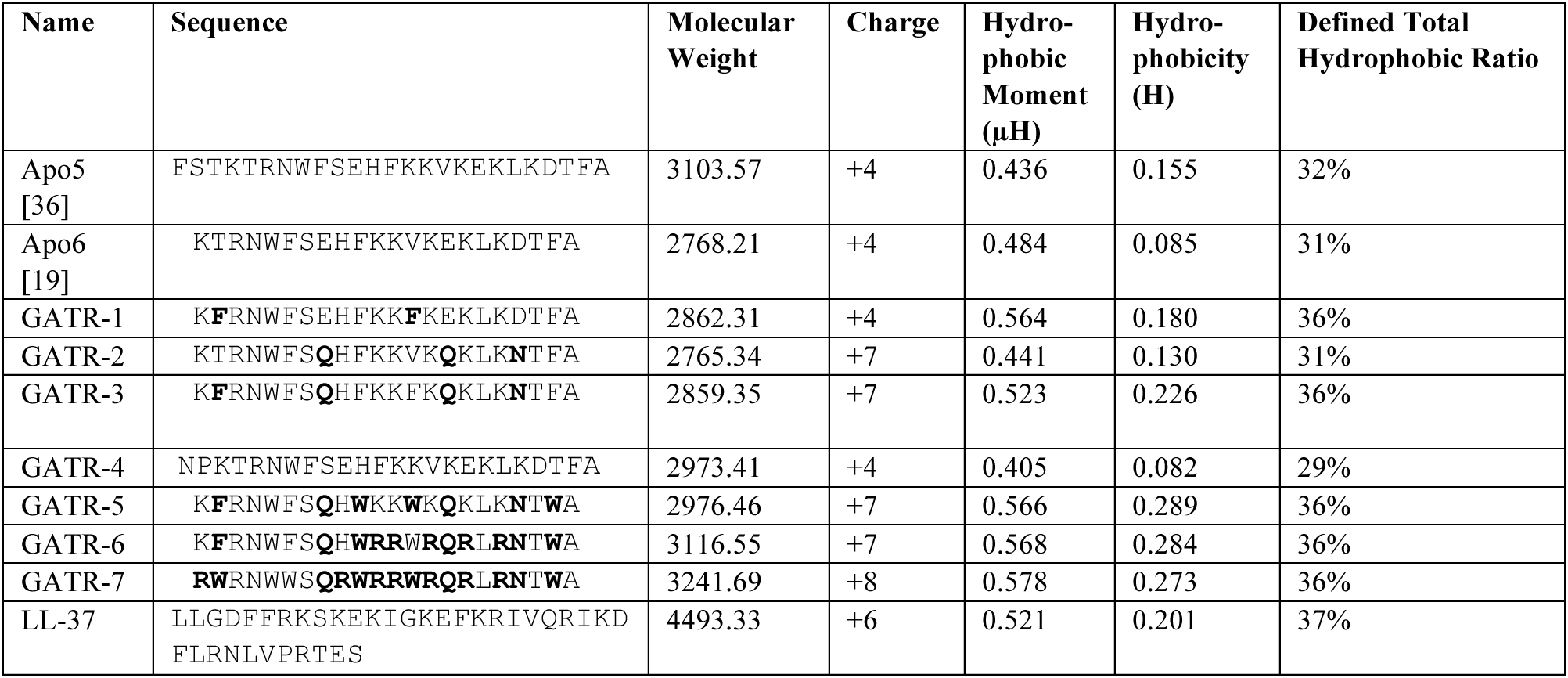
Sequences and physico-chemical properties of GATR antimicrobial peptides. Physico-chemical properties and helical wheels were calculated using Heliquest [21] and APD3 [22].

GATR-1 was produced by replacing the native threonine (T) in position 2 with a phenylalanine (F), and substituting valine (V) at position 13 with phenylalanine (F). These changes increase the hydrophobic moment of the helical peptide from 0.484 µH for Apo6 to 0.564 µH as well as raising hydrophobicity from 0.085 H to 0.180 H.

GATR-2 was produced by replacing glutamic acid (E) at position 8 with glutamine (Q), glutamic acid (E) at position 15 with glutamine (Q), and aspartic acid (D) at position 19 with asparagine (N). These alterations to the sequence raise the peptides positive charge from +4 (Apo6) to +7 and hydrophobicity to 0.130. However, these changes also reduce the hydrophobic moment to 0.441 µH.

GATR-3 combines the T2/V13 and E8/E15/D19 amino acid substitutions of GATR-1 and GATR-2. These sequence modifications increase the overall peptide charge to +7, hydrophobic moment to 0.523 µH, and net hydrophobicity to 0.226.

GATR-4 was produced by adding NP to the N-terminus because N-capping peptides, particularly with a proline residue, has been reported to increase peptide stability and decrease protease susceptibility [34]. Add numbers about charge, HM and H from table.

GATR-5 was produced by combining the GATR-2 alterations with substitutions of phenylalanine (F) at position 10 to tryptophan (W), valine at position 13 to tryptophan (W) and phenylalanine (F) at position 21 to tryptophan (W). These modifications increase the peptide charge from +4 (Apo6) to +7, the hydrophobic moment to 0.566 µH, and hydrophobicity to 0.289 H.

In GATR-6, the sequence of GATR-5 has been further modified by replacing the lysine (K) residues K11, K12, K14, K16, and K18 with arginine (R) residues. The physicochemical properties of GATR-6 are nearly identical to those of GATR-5 and also GATR-3. Both GATR-5 and GATR-6 have a net charge of +7, hydrophobic moments of 0.566 µH and 0.568 µH respectively, and hydrophobicities of 0.289 H and 0.284 H respectively.

GATR-7 was produced from the GATR-6 sequence by substituting the lysine (K) at position 1 with arginine (R), the phenylalanine (F) at position 2 to tryptophan (W), the phenylalanine (F) at position 6 to tryptophan (W) and the histidine (H) at position 9 to arginine (R). Due to these substitutions, GATR-7 is predicted to have a net charge of +8, which is higher than that of the other GATR peptides. Additionally, the hydrophobic moment of GATR-7 is 0.578 µH and its net hydrophobicity is calculated to be 0.273 H. These values are similar to those calculated for GATR-3, GATR-5 and GATR-6.

### GATR peptides are antibacterial against *Francisella tularensis*

Apo6 has been shown to have activity against a broad range of pathogens (*Bacillus cereus*, *Escherichia coli*, *Pseudomonas aeruginosa*, and *Staphylococcus aureus*, as well as drug-resistant strains of *E. coli, P. aeruginosa, S. aureus*, and *Acinetobacter baumannii*) in low salt buffer (EC_50_) [16, 19]. Apo6 shares a salt-sensitive phenotype with LL-37 [14, 35] and was found to be inactive in Muller-Hinton broth against these bacteria in MIC assays [16, 19]. We first tested Apo6 and the GATR peptides in MIC assays against *F. tularensis* LVS. Similar to its activity against other bacteria, Apo6 had no observable MIC against *F. tularensis* LVS at the concentrations tested. In addition, GATR-1, GATR-2, GATR-3, and GATR-4 were found to be inactive under these conditions; however, some inhibitory activity was observed when GATR-5, GATR-6, and GATR-7 are tested, with 85% inhibition at 41.5 µg/ml in the case of GATR-7 (**Figure 1).** It appears that the antibacterial activity in media increases along with the hydrophobic moment.

**Figure 1.**
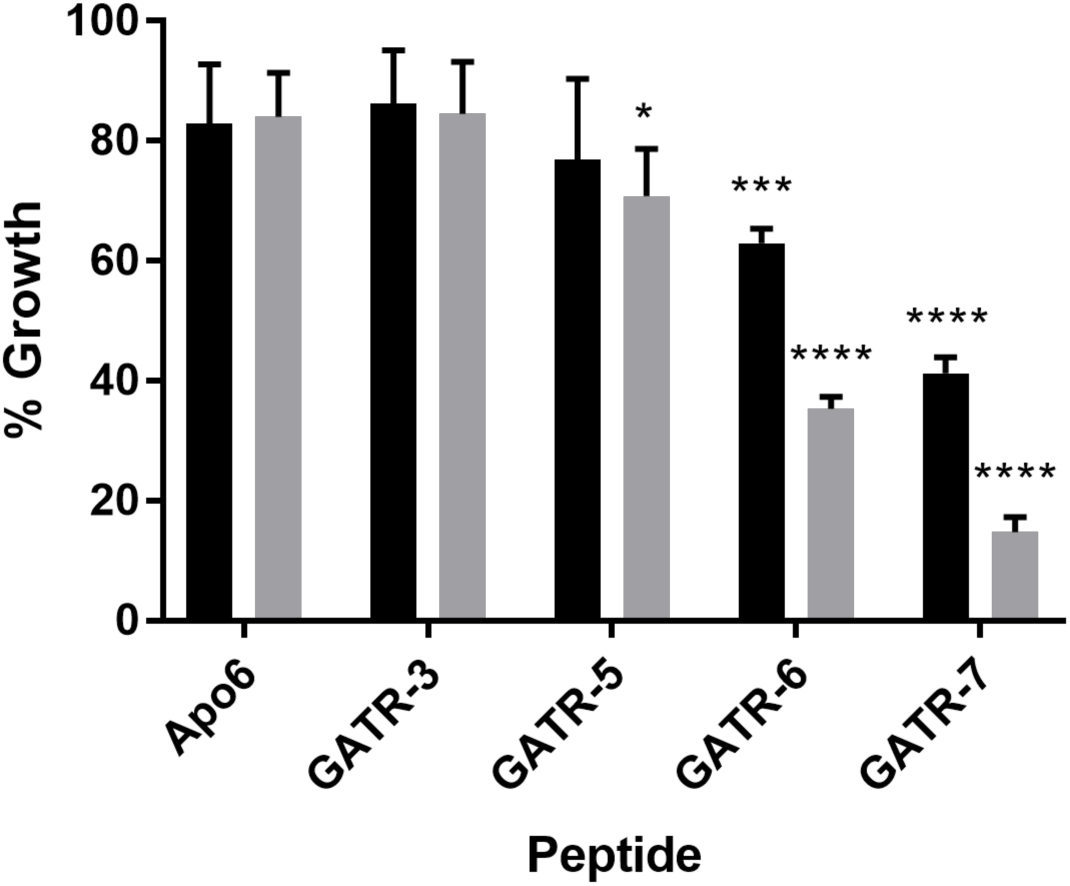
GATR peptides have improved antibacterial activity in broth against *F. tularensis* LVS. MIC assays were performed in Cation-adjusted Mueller Hinton Broth with 2% IsoVitaleX (black=21 µg/ml; gray=42 µg/ml) on *F. tularensis* LVS with 5 replicates per experiment. Experiment was performed twice. Results were analyzed using a 2way ANOVA with Sidak’s multiple comparison against Apo6. Error bars indicate standard deviation. (* p<0.05; *** p<0.001; ****p<0.0001).

Next, we tested the antimicrobial activity of Apo6 and its derivatives against *F. tularensis* LVS and *F. tularensis* NIH B38 strain in a low salt buffer, which is an alternate measure of antimicrobial activity reported as EC_50_ values, shown in **Table 2** [16, 19, 26, 27, 34-36]. We performed these experiments using LL-37 as a positive control, which was found to be highly effective against *F. tularensis* LVS (EC_50_= 0.209 µg/ml), similar to the EC_50_ reported for *F. novicida* [27]. As shown in **Table 2**, it was found that the EC_50_ values of the GATR peptides were generally lower than that of Apo6, which has an EC_50_ value of 6.82 µg/ml against *F. tularensis* LVS and 16.3 µg/ml and *F. tularensis* NIH B38, with the exception of GATR-4, which had a similar EC_50_ against *F. tularensis* LVS (11.0 µg/ml) but was not effective at concentrations tested against *F. tularensis* NIH B38. Four peptides (GATR-3, GATR-5, GATR-6, and GATR-7) had EC_50_ values lower than 2 µg/ml against both strains of *F. tularensis*, and thus were selected as the most effective peptides against *Francisella.* For comparison, the EC_50_ of levofloxacin for *F. tularensis* LVS is 0.00827 µg/ml (8.27 ng/ml) [36].

Apo6, the four most effective GATR variants, and LL-37 were then tested against the highly virulent strain *F. tularensis* SchuS4 in MIC and low salt assays. Results are shown in **Table 3**. Although the parent peptide Apo6 is not effective against *F. tularensis* SchuS4, GATR-3, GATR-6, and GATR-7 are each moderately effective against this strain, with EC_50_ values around 30 µg/ml. Interestingly, while these 4 peptides are not particularly effective against less virulent strains of *F. tularensis* in MIC assays, GATR-7 displays comparatively strong activity with a MIC of 41.7 µg/ml.

**Table 3.** Antibacterial activity of selected GATR peptides against F. tularensis tularensis (Ftt) SchuS4. The more active GATR derivatives were tested against *Ftt* SchuS4 in CAMHB with 2% IsovitaleX and in Buffer Q. For statistical comparison, the 95% confidence intervals (p < 0.05) are listed. The EC50 values are also expressed as µM for direct comparison.

| Peptide Name | MIC [µg/ml] | EC<sub>50</sub> (95% CI) [µg/ml] | EC<sub>50</sub> [µM] |
| --- | --- | --- | --- |
| Apo6 | Not tested | Not active | Not active |
| GATR-3 | >83.3 | 28.6 (13.3 to 61.6) | 10.0 |
| GATR-6 | >83.3 | 32.3 (15.8 to 66.2) | 10.4 |
| GATR-7 | 41.7 | 24.2 (wide) | 7.47 |
| LL-37 | Not tested | 0.562 (0.195 to 1.61) | 0.125 |

### GATR peptides interact with the cytoplasmic membrane of *F. tularensis* LVS

As part of their mechanism of action, antimicrobial peptides can cause bacterial membrane disruption, ranging from slow leakage of cellular contents owing to membrane thinning to formation of large monomeric pores that can lead to cell death [37]. In order to evaluate the interaction between the peptides and the bacterial cytoplasmic membrane, we conducted two fluorescence-based studies. One of the ways in which the structural integrity of cell membrane can be compromised is through disruption of membrane potential [38]. We assessed the depolarization of bacterial membranes using DiSC_3_(5), a membrane potential sensitive dye, which intercalates itself in the lipid bilayer resulting in the self-quenching of the dye [38]. If depolarizing compounds are added, the potential decreases, and DiSC_3_(5) is released into the solution causing an increase in fluorescence relative to the reduction of membrane potential [31, 37, 38]. **Figure 2A** indicates a concentration-dependent increase in fluorescence when *F. tularensis* LVS was treated with two different concentrations of peptides (10 µg/ml and 1 µg/ml). Apo6 and the GATR peptides dissipated the membrane potential in *F. tularensis* LVS at 1 µg/ml, indicating that depolarization of cytoplasmic membrane is a primary mechanism of action of Apo6 and its derivatives. In addition, the derivatives were much more effective in disrupting the membrane potential at 10 µg/ml compared to the parent peptide Apo6 (p values < 0.0001).

**Figure 2.**
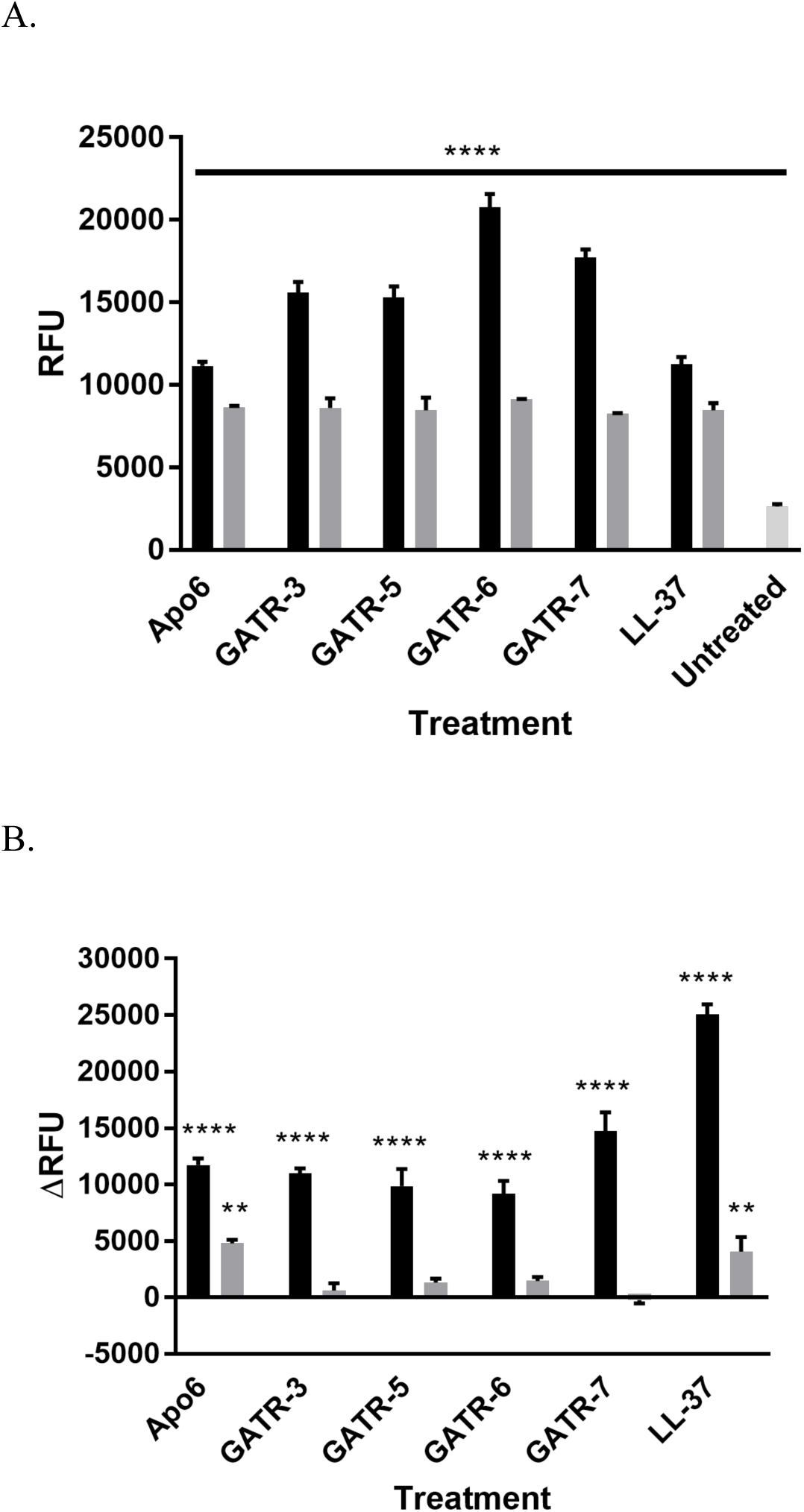
GATR peptides disrupt the bacterial membrane of *F. tularensis* LVS. A. Membrane depolarization was measured using DiSC3(5) in 10 mM phosphate buffer with at least 2 replicates per experiment (black=10 µg/ml; gray=1 µg/ml). Experiment was performed 3 times. B. Pore formation or greater membrane perturbation was measured using ethidium bromide in 10 mM phosphate buffer with 3 replicates per experiment (black=10 µg/ml; gray=1 µg/ml). Experiment was performed 3 times. Results were analyzed using a 1way ANOVA with Dunnet’s multiple comparisons. Error bars indicate experimental variation. (** p<0.01; **** p<0.0001)

Greater disruption can lead to the formation of larger, less transient holes or pores in the bacterial membrane, which will lead to bacterial death. To examine this effect, we conducted a membrane disruption assay using ethidium bromide (EtBr). This larger molecule will pass through a damaged membrane and intercalate with the bacterial DNA resulting in increased fluorescence proportional to the level of membrane disruption. We observed that *F. tularensis* LVS was sensitive to pore-formation by Apo6 and its derivatives (Figure 2B), evidenced by a significant RFU difference between the control and treated bacteria (p-values <0.05). At 10 µg/ml, all peptides except GATR-7 demonstrate a significant change in RFU, indicating pore-formation by most of these peptides. Apo6 and GATR-5 also display significant pore formation at a lower concentration of 1 µg/ml. However, GATR-3, GATR-6, and GATR-7 do not show significant pore formation compared to the untreated bacteria at 1 µg/ml. LL-37 was used as positive control in the depolarization and pore formation studies of the peptides [19, 28].

### GATR peptides bind *F. tularensis* LVS lipopolysaccharide (LPS)

Lipopolysaccharide (LPS) is a major structural component of the Gram-negative bacterial outer membrane and protects bacteria from antimicrobial compounds [11, 30]. LPS from *E. coli* and other gram-negative bacteria is the endotoxin and activates innate immunity through binding TLR4 receptors [39]. The overall positive charge on cationic antimicrobial peptides assists them to form strong electrostatic interactions with the negatively charged LPS in the membrane of Gram-negative bacteria neutralizing the overall negative charge [40, 41]. The binding of cationic antimicrobial peptides with LPS of Gram-negative bacteria has a major effect on the stability of bacterial membranes. It has been previously demonstrated that several cationic antimicrobial peptides including LL-37, SMAP-29, and CAP18 can bind LPS [42–44]. Some cationic antimicrobial peptides have been shown to reduce the host immune response to LPS by binding and sequestering it [42].

*Francisella* LPS is unusual among gram-negative LPS as it does not induce a strong pyogenic response or activate TLR4 signaling [45–47]. Cationic antimicrobial peptide-LPS binding could lead to greater interaction between the cationic antimicrobial peptides and the *Francisella* bacterial membrane, which might enhance the activity of the peptide against the bacteria. Thus we decided to investigate the ability of Apo6 and GATR peptides to bind purified LPS from *F. tularensis* LVS. In order to analyze the binding between peptides and LPS, we employed a DMMB dye LPS-binding assay. The positively charged dye competes with the positively charged peptide to bind to the negatively charged moieties on the LPS. Upon binding, the dye changes color from blue to purple/pink. As shown in **Figure 3**, we found that although the parent peptide Apo6 does not significantly bind *F. tularensis* LVS LPS, the GATR peptides tested significantly bind this LPS (p value <0.05). GATR peptides with greater charge and hydrophobicity (GATR-6 and GATR-7) bind this LPS in greater amounts than do less charged and hydrophobic peptides (GATR-3 and GATR-5). Thus, LPS binding might contribute to the anti-*Francisella* mechanism of the GATR peptides.

**Figure 3.**
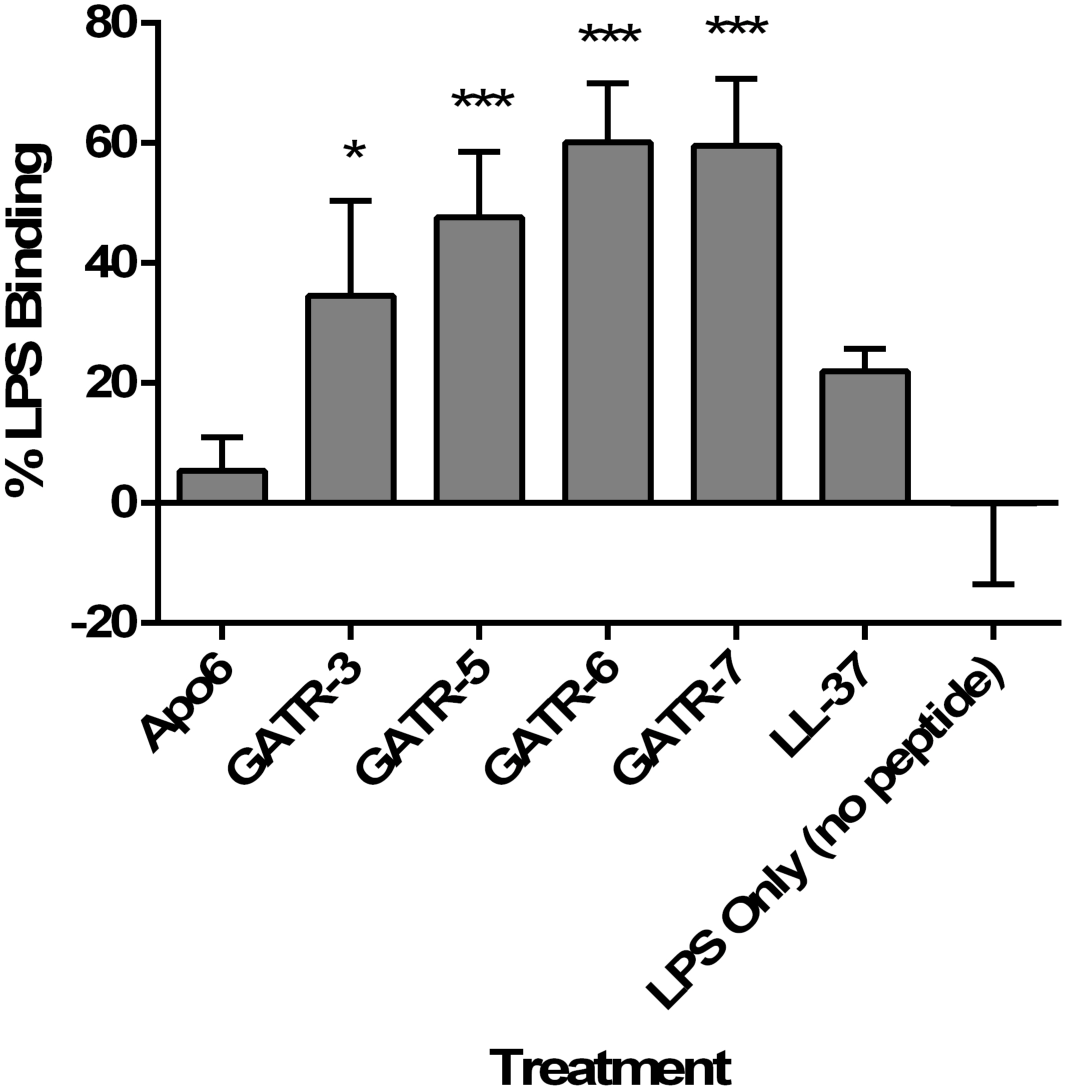
GATR peptides bind *F. tularensis* LVS LPS. 150 µg/ml of LPS was incubated with 10 µg/ml of peptide in distilled endotoxin-free water for 1 h and then added to DMMB. The experiment was performed twice with 3 replicates per experiment. Results were analyzed using a 1way ANOVA with Dunnett’s multiple comparisons tests. (* p<0.05; ** p<0.01; *** p<0.001) Error bars indicate standard deviation.

### Toxicity of the GATR peptides

We sought to further down-select the peptides by testing for potential toxicity. To examine whether the GATR peptides may be toxic to mammalian cells (particularly those peptides with higher charge), we performed hemolysis assays, cytotoxicity assays using the MTT assay, and toxicity experiments in *G. mellonella* waxworms. First, hemolysis assays using sheep red blood cells were performed at peptide concentrations of 100 µg/ml for 1 h [19, 31, 34]. GATR-5, −6, and −7 showed statistically significant hemolysis, in particular GATR-6 and GATR-7, each of which had hemolysis levels of greater than 20% of RBCs (**Figure 4A**). Next, we measured cytotoxicity of the GATR peptides by using the MTT assay as a measure of cell viability following peptide treatment [19, 31, 34]. A549 human lung epithelial cells and HepG2 liver cells were treated with 100 µg/ml peptide for 24 h. Shown in **Figures 4B** and **4C**, some statistically significant suppression of cell proliferation was seen in A549 cells for GATR-3, GATR-6, and GATR-7; however, no peptides show statistical suppression of cell growth when tested against HepG2 cells.

**Figure 4.**
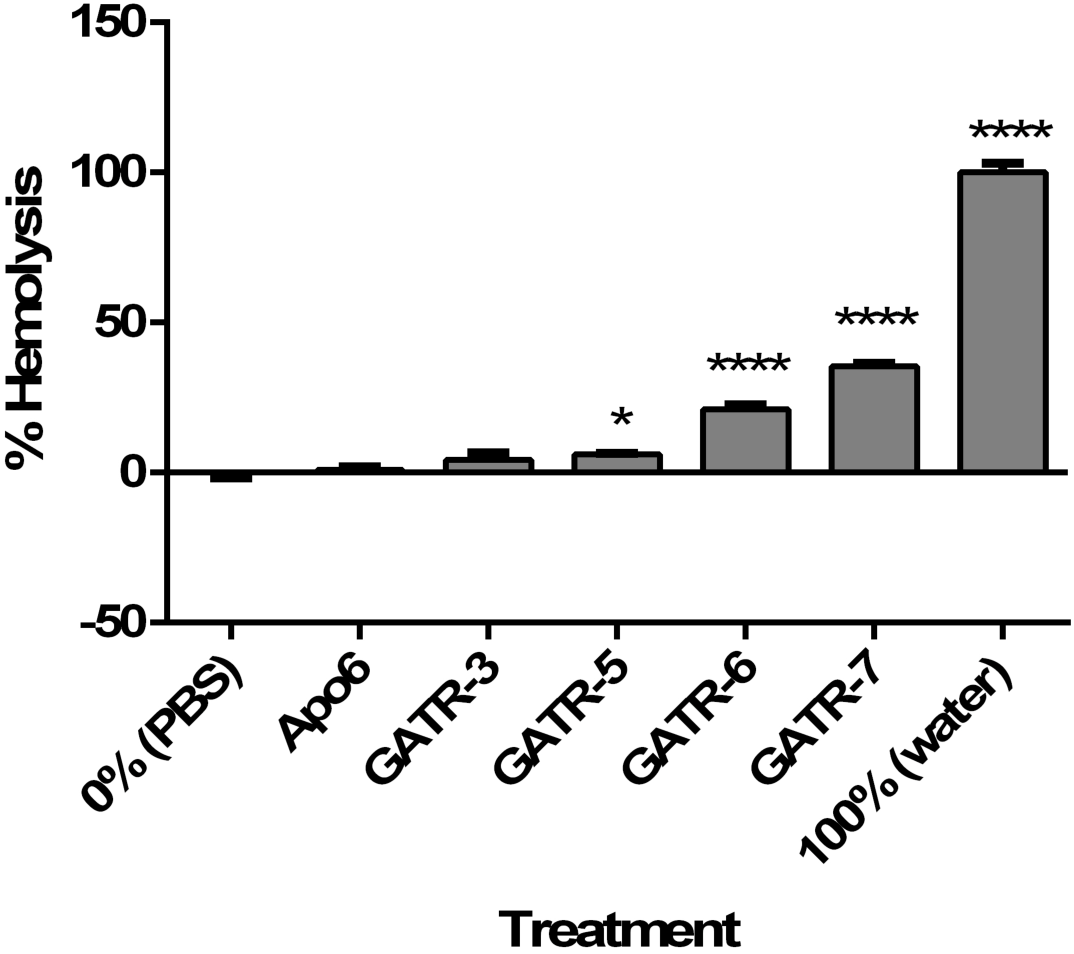
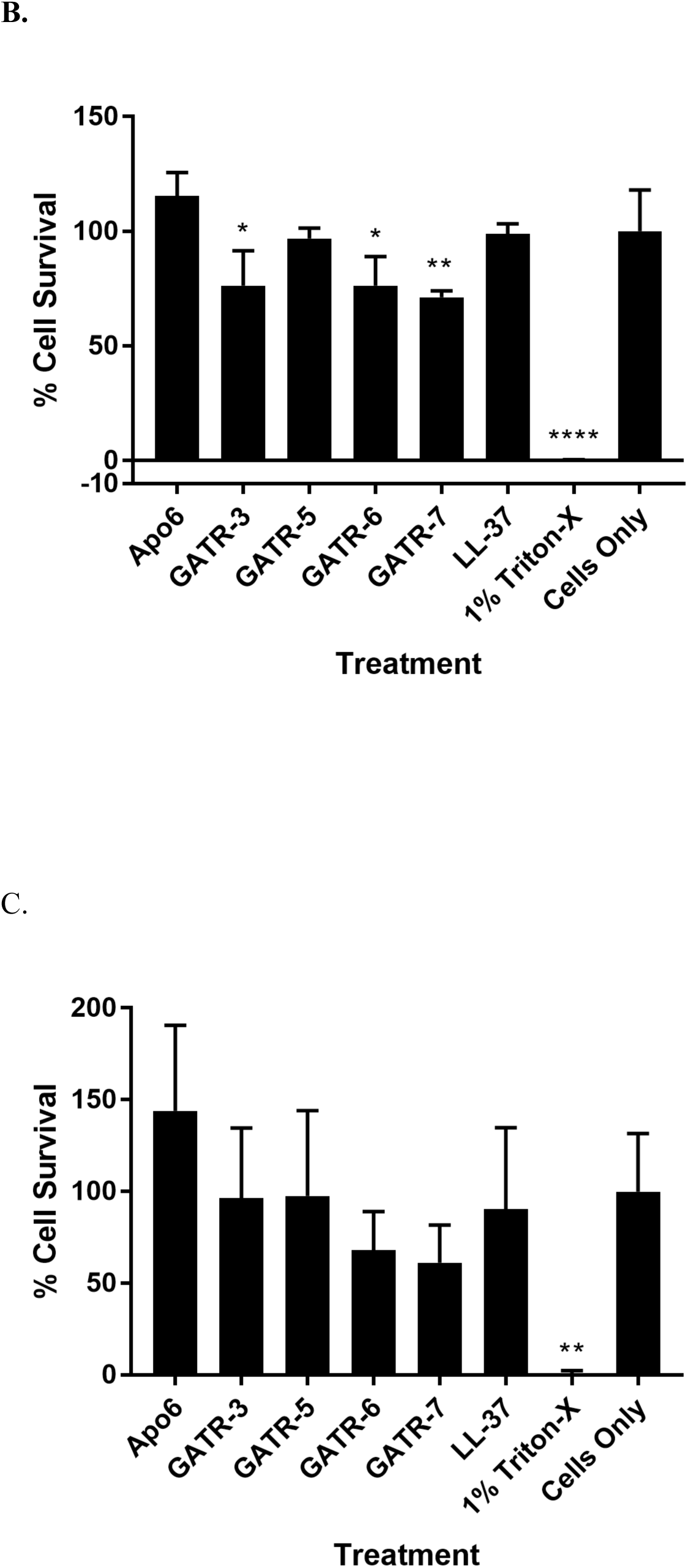
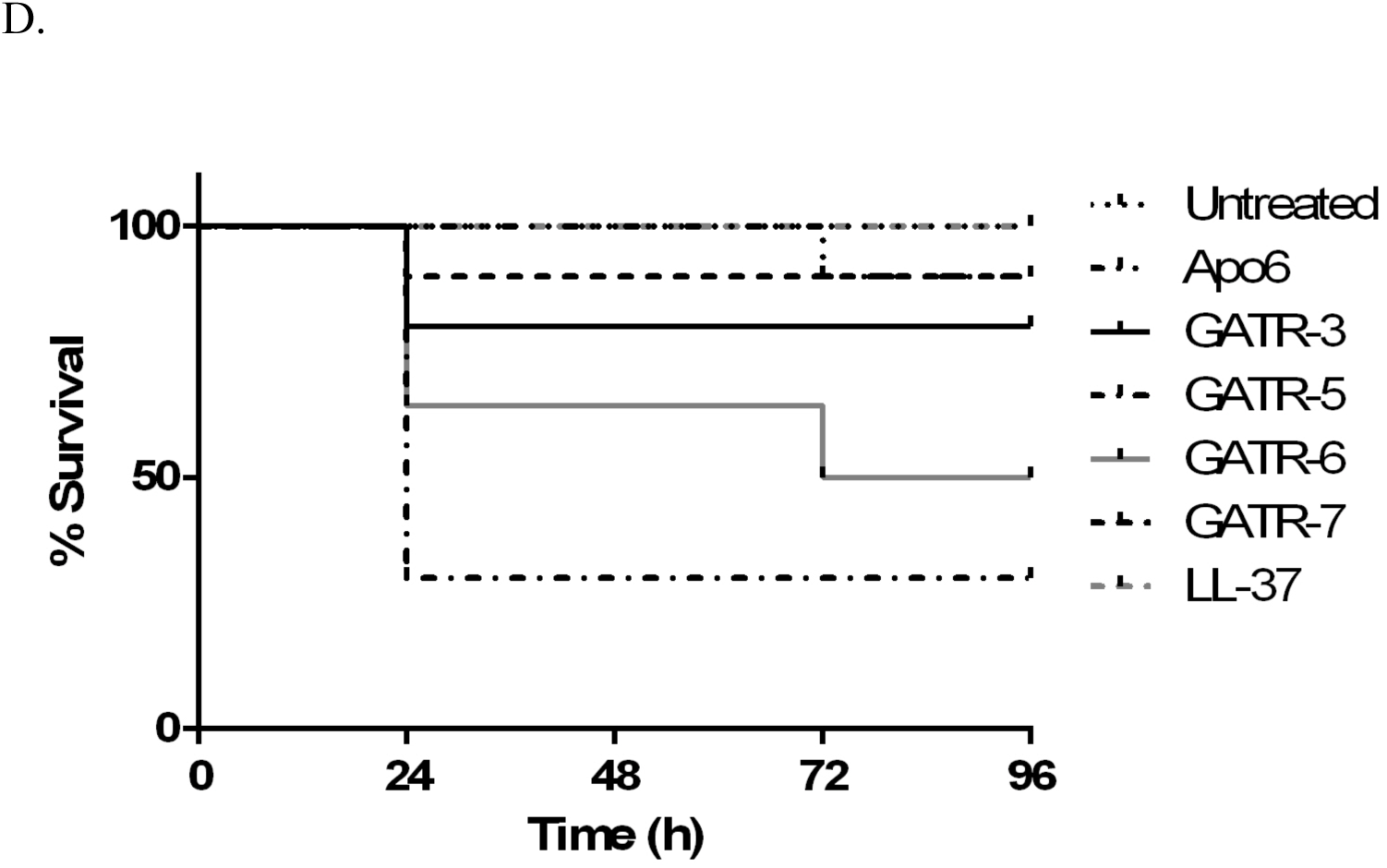
Toxicity of GATR peptides. **A.** Hemolysis assay using 2% sheep red blood cells. Peptides are reconstituted in sterile PBS. For 0% hemolysis, RBCs are exposed to PBS. For 100% hemolysis, RBCs are exposed to sterile water. Experiment was performed twice with 6 replicates per experiment. Results were analyzed using a 1way ANOVA with Dunnett’s multiple comparisons. **B.** MTT cell proliferation assays using A549 human lung epithelial cells and **C.** HepG2 human hepatocytes with 24 h exposure to 100 µg/ml peptide. Experiments were performed twice each with 3 replicates per experiment. Results were analyzed using a 1way ANOVA with Dunnett’s multiple comparisons. (* p<0.05; ** p<0.01; **** p<0.0001) **D.** Toxicity in *G. mellonella* larvae was measured by injecting each worm with 10 µg of peptide (10 larvae/group). Survival was measured for 48 h.

Toxicity assays were also performed in the *G. mellonella* waxworm model. In groups of 10, each larvae received 10 µg of peptide, and survival was assessed for 48 h. After this time period, waxworms treated with GATR-3, GATR-6, and GATR-7 were not found to have significant death as measured by larvae survival (**Figure 4D**). However, GATR-5 treated waxworms had only 30% survival, indicating that this peptide could potentially be toxic in an animal model (p=0.0014), and so this peptide will not be carried forward to the *in vivo* testing.

### Waxworm *in vivo* infection survival assay

Analysis of the efficacy of antimicrobials utilizing *in vivo* models is conducted to assess the anti-infective potential of the drug in an infected animal. Ideally a mammalian animal model should be employed in order to test the *in vivo* capabilities of antimicrobials; however, alternative models may be appropriate for screening of lead antimicrobial candidates (EC_50_ activity ≤10 µg/ml). *Galleria mellonella*, the greater wax moth, has been proposed as an alternative model that is relatively easy to obtain and has a system of antimicrobial protection similar to that of mammals. These factors make larvae of *G. mellonella* a model of infection for various pathogenic microorganisms [2, 32, 48, 49]. *G. mellonella* has been previously used as an infection model for *in vivo* effect of antimicrobial peptides and antibiotics against *Francisella spp.* infections [2, 48].

In the current study, to evaluate the ability of selected antimicrobial peptides to prolong survival of infected *G. mellonella*, larvae were infected with *F. tularensis* LVS and then treated with a single dose of 10 ng of peptides. Shown in **Figure 5**, *G. mellonella* showed statistically significant improved survival when compared to untreated groups (p<0.05) when treated with Apo6 and GATR peptides, with GATR-3 having the strongest effect (80% survival, p=0.0001). The parent peptide, Apo6, was the next best candidate (60% survival, p=0.0002). While GATR- 5-treated *G. mellonella* initially demonstrated a strong prolonged survival rate, all of the larvae succumbed to infection by 120 h (0% survival, p=0.0008). GATR-6- (30% survival, p=0.045) and GATR-7- (30% survival p value= 0.0015) treated waxworms also showed significant prolonged survival.

**Figure 5.**
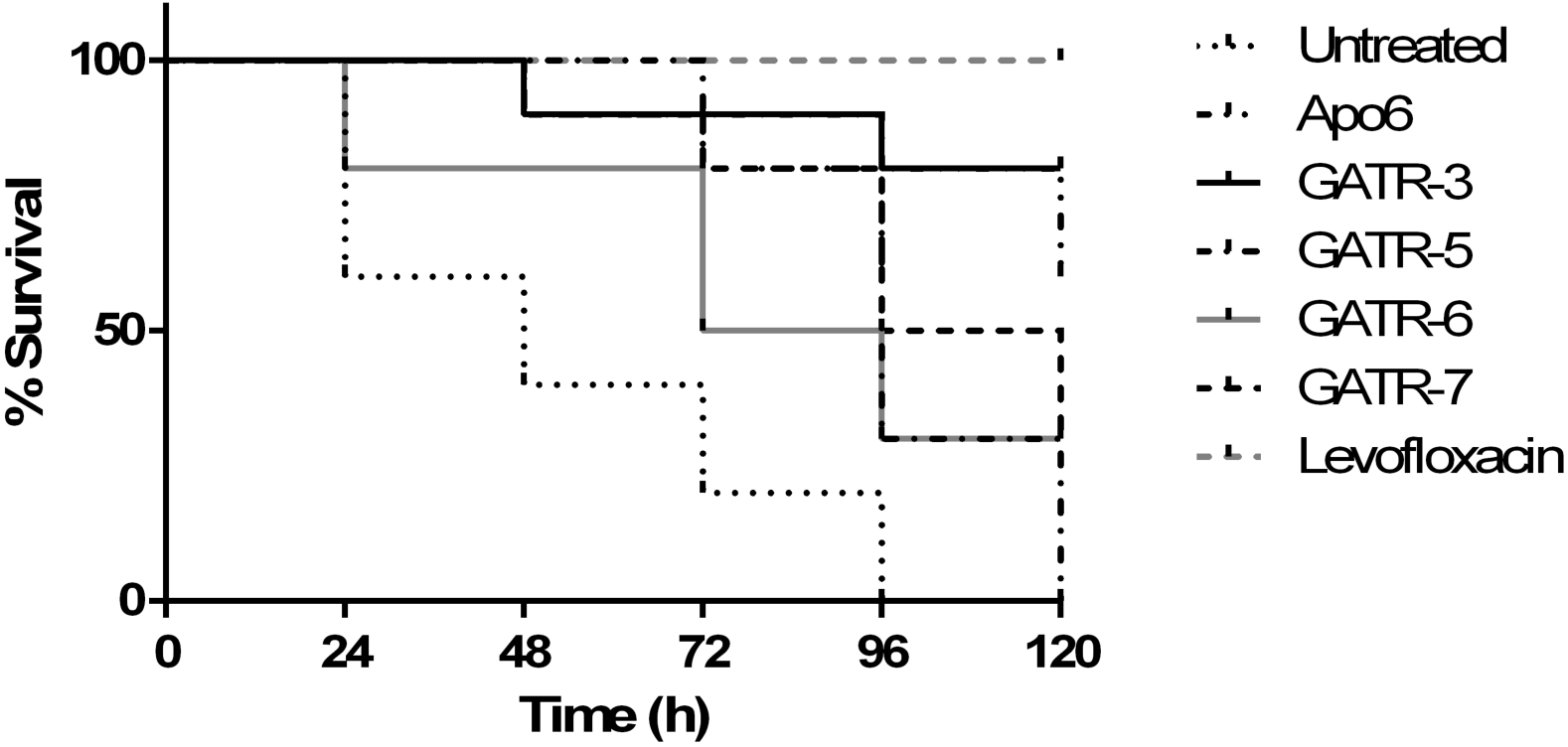
*G. mellonella* survival following GATR peptide treatment. *G. mellonella* larvae were infected with *F. tularensis* LVS and treated with a single injection of 10 ng peptide or 10 µg levofloxacin (10 larvae/group). Survival was monitored for 120 h after infection.

### Murine model *in vivo*

After down-selecting our lead peptides through the *G. mellonella* invertebrate model, the 3 lead peptides were tested in a murine model of pulmonary tularemia. GATR-3, GATR-6, and GATR-7 were tested. In addition, we tested Apo6, LL-37, and D-LL-37 in this model. LL-37 had stronger activity *in vitro* than any of the GATR peptides. Previously, Flick-Smith et al evaluated the use of LL-37 as a post-exposure intranasal therapy in the treatment of pulmonary tularemia by delivering the peptide directly to the lungs [20]. In that report, LL-37 extended mean time till death but did not increase survival in treated animals. In our study, systemic peptide delivery via the intra-peritoneal (IP) route of treatment appears to have no adverse effect on survival in response to this peptide, i.e. the peptide was not toxic when delivered systemically. Previously, we have found that D-LL-37 has increased activity antimicrobial and protease resistance [35, 36], making it an attractive peptide to use *in vivo*. However, in our study, D-LL-37 had similar activity in this animal model to LL-37 in that it did not extend survival or rescue mice, nor did these peptides lessen signs of disease. Survival data and health scores are shown in **Figure 6A** and **7B**.

**Figure 6.**
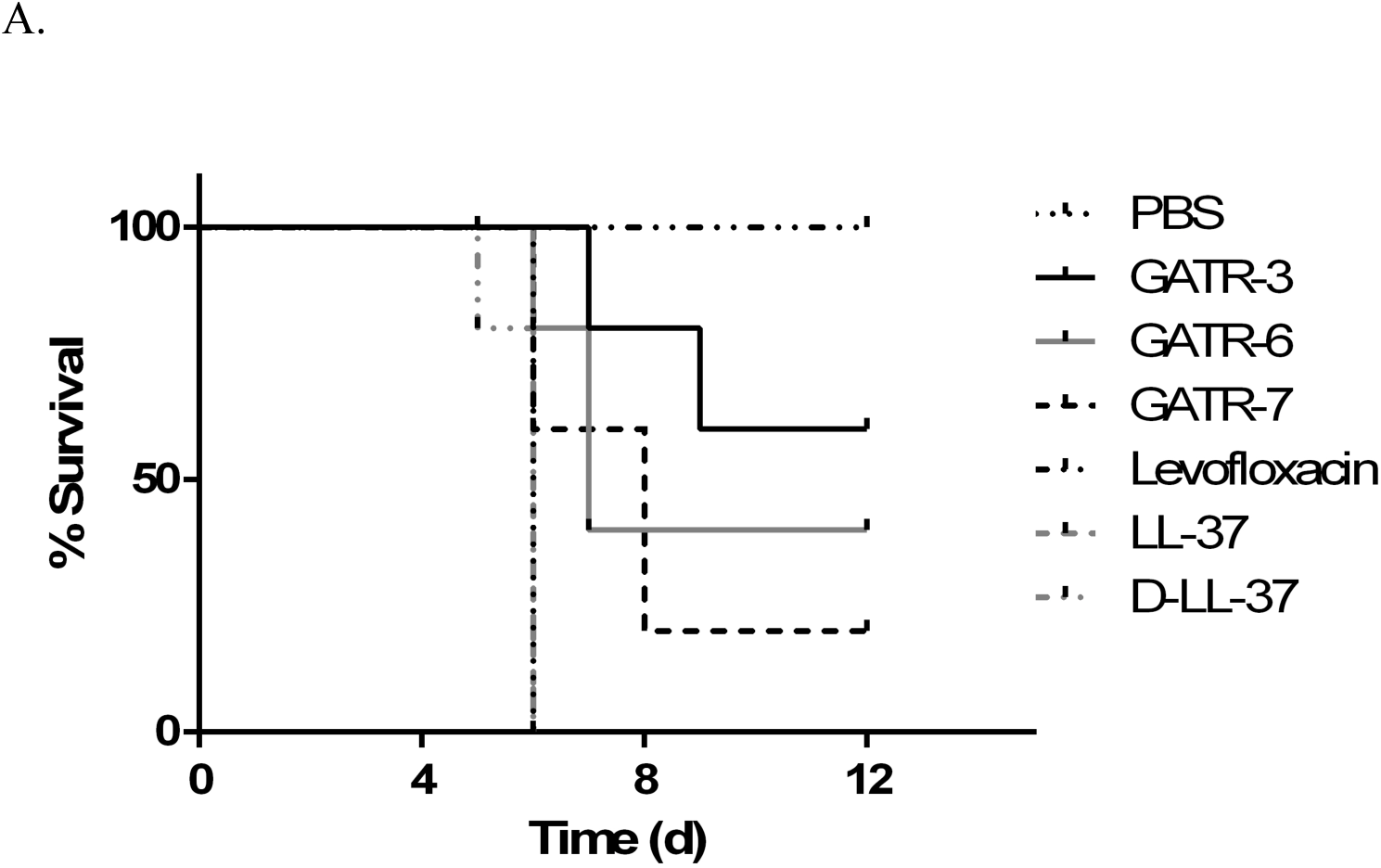
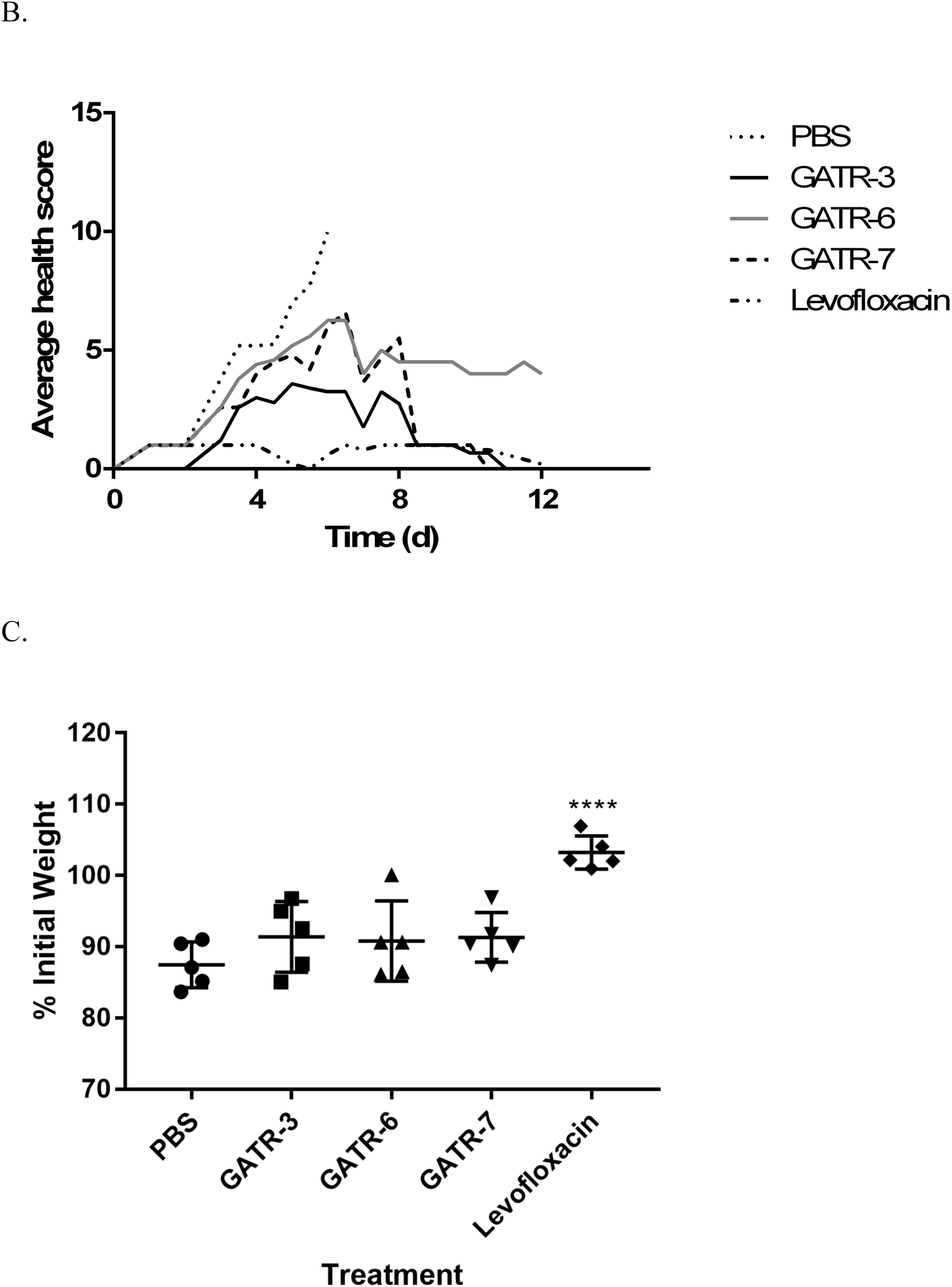
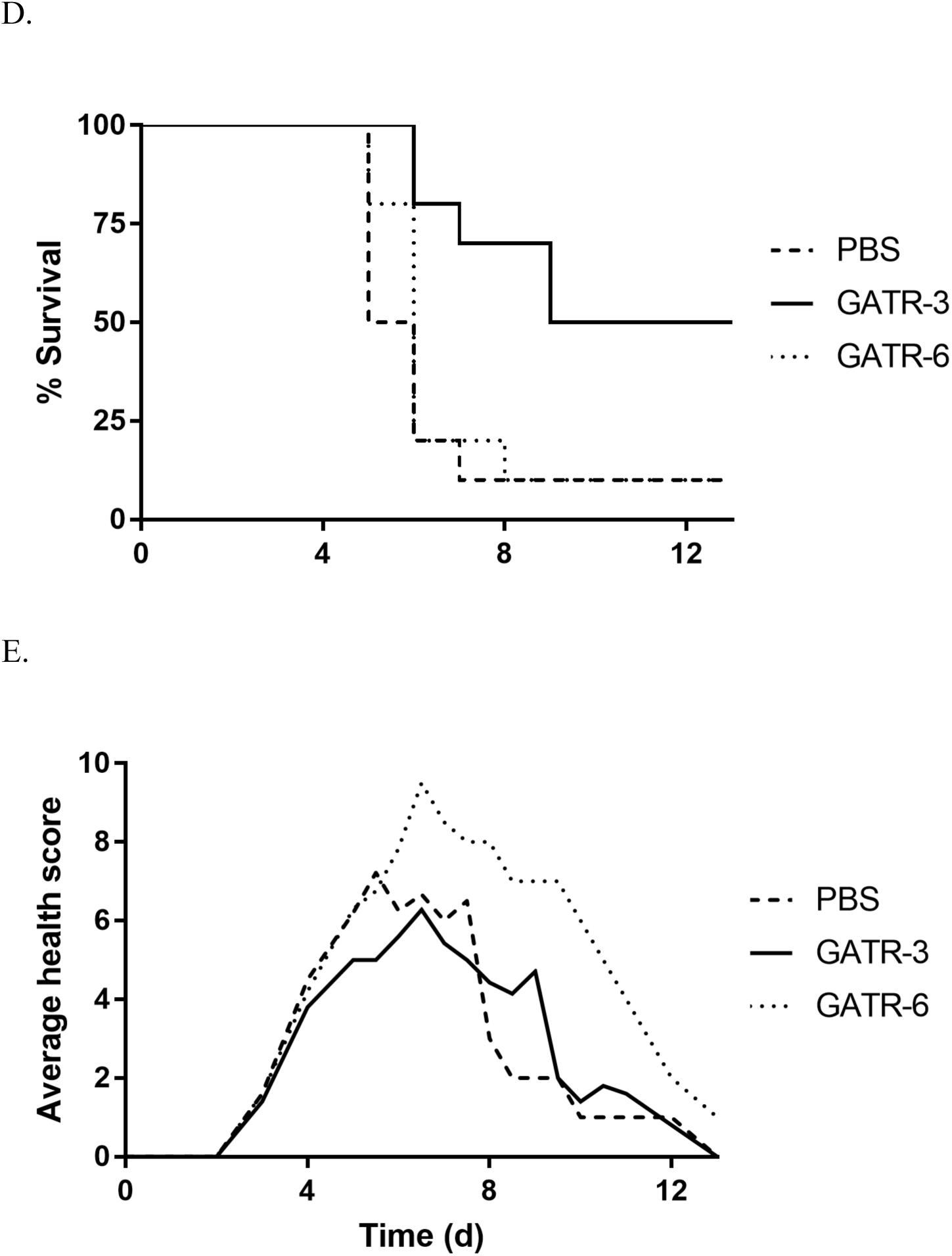
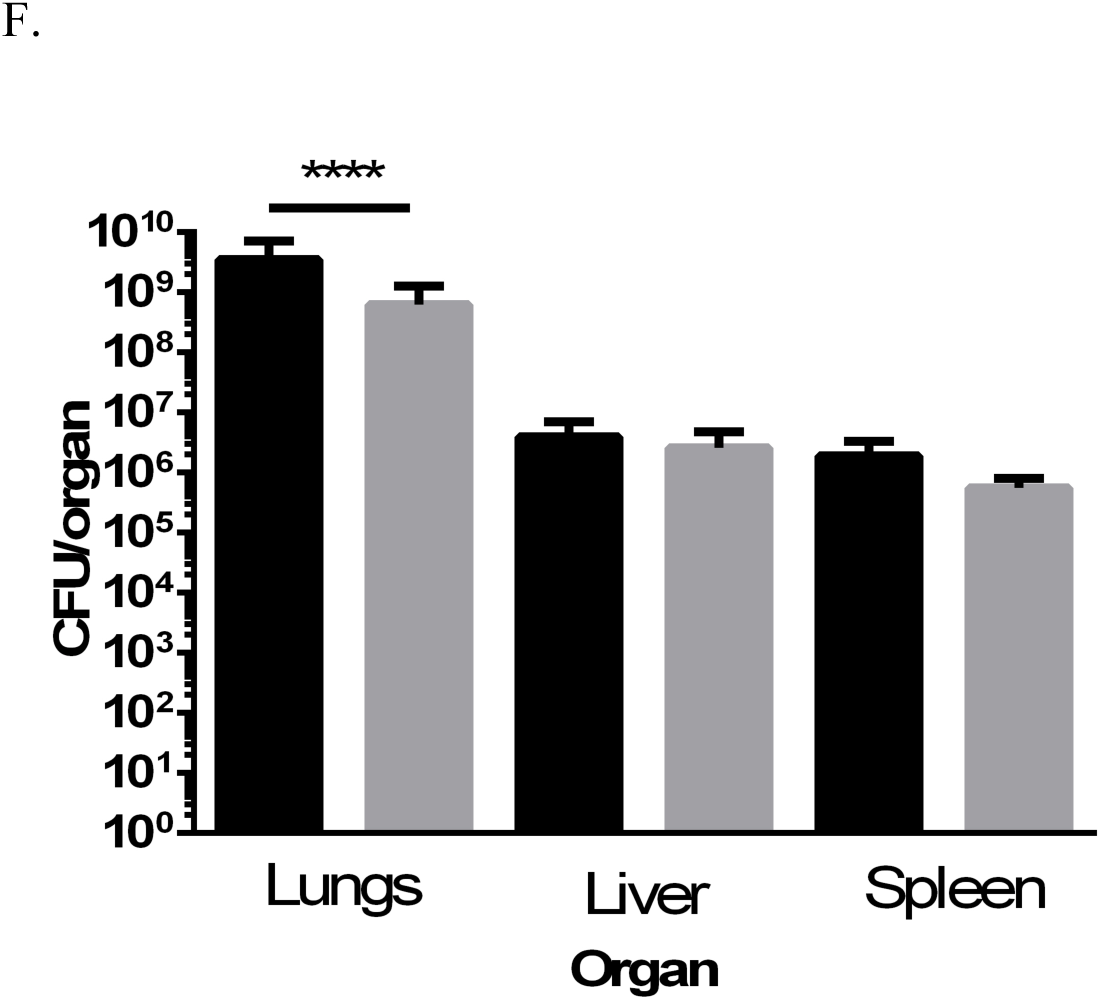
GATR Peptide treatment of *F. tularensis* LVS infected mice. BALB/c mice were infected with 50 LD50 of *F. tularensis* LVS and treated with peptide 24 h before and 3, 24, and 48 h after infection (5 mice/group). A. Survival curves of mice with prophylactic treatment, B. Average health scores over course of study, C. percent initial weight on day 4 after infection, in which results were analyzed using a 1way ANOVA with Tukey’s multiple comparisons (**** p<0.0001) Next, BALB/c mice were infected with 10 LD50 of *F. tularensis* LVS and treated with peptide 3, 24, and 48 h after infection (5 mice/group). D. Survival curves of mice with post-infection treatment only E. Average health scores of mice during survival study. F. Organ burden study comparing untreated (black bars) and GATR-3-treated (gray bars) organs (3 mice/group). Lungs, liver, and spleen were harvested on day 4 after infection homogenized in PBS, and plated on chocolate agar. Results were analyzed using a 1way ANOVA with Tukey’s multiple comparisons. (**** p<0.0001)

In the first set of experiments, mice (5/group) were given 1 prophylactic peptide treatment 1 day before infection (Day −1) and 3 treatments after infection at 3, 24, and 48 h. Each intra-peritoneal (IP) injection delivered 5 mg/kg of peptide. Survival is shown in **Figure 6A**. In this model, GATR-3 was found to be the most successful peptide, with 60% survival (p=0.0047) compared to the PBS-injected control, which had a mean time till death (MTD) of 6 days and 0% survival. GATR-6 saved 40% of mice (p=0.0237) compared to the control. GATR-7 saved 20% of mice, but Mantel-Cox tests indicate this was not significant (p=0.0736). When average health scores were examined in **Figure 6B**, GATR-3 delayed the time of disease onset from day 1 to day 3 and lessened severity of clinical signs over the course of infection. GATR-6 and GATR-7 did not delay disease onset, but severity of signs were slightly lessened compared to the untreated control. For this set of experiments, mice were weighed daily in the morning. Weights from day 4 are compared in **Figure 6C**. None of the GATR peptide treated mice show a significant difference in weight change compared to the untreated control.

In the second set of experiments, the prophylactic pre-treatment was not given and larger groups were used (10 mice/group). Mice received only the 3 treatments after infection at 3, 24, and 48 h. Because LL-37, D-LL-37, and GATR-7 were not found to significantly rescue mice in the first set of studies, they were not used in these experiments. Only GATR-3 and GATR-6 were tested. In the survival study shown in **Figure 6D**, GATR-3 once again was the most successful peptide, with 50% survival (p=0.0053) compared to the untreated control, which had 10% survival and a mean time till death of 5.5 days. However, GATR-6 was not found to have significant survival in this study (10% survival, p=0.4347), though mean time till death (MTD) was extended to 6 days. This may indicate the prophylactic treatment was important for the efficacy of GATR-6. When average health scores were compared in **Figure 6E**, differences between peptide-treated and untreated groups were not as apparent as in **Figure 6B**. All groups had signs of disease onset on day 3. Disease signs of GATR-3-treated mice were only slightly lessened compared to the untreated group, while signs of GATR-6-treated mice were slightly more severe than the untreated mice.

Because GATR-3 rescued mice in both sets of experiments, an organ burden study was performed with this peptide on day 4 to determine the bacterial burden in the lungs, spleen, and liver (3 mice/group), as shown in **Figure 6F**. Though no significant difference was found between the bacterial burden in the spleens and livers of GATR-3-treated and untreated mice, the bacterial numbers were found to be significantly lower in lungs (p<0.0001). Thus, the survival benefit may have been due to reduced lung burden as opposed to overall bacterial clearance.

## Discussion

*Francisella* is highly resistant to cationic cyclic peptide antibiotics such as polymyxin B. Indeed, *Francisella* selective growth media contains 100 mg/ml polymyxin B [50]. The resistance to polymyxin B is thought to be due to the special structure of the lipopolysaccharide (LPS) of *Francisella* [46, 51]. Thus, *Francisella* is considered to be resistant to this class of cyclic peptide antibiotics, which are sometimes called cationic antimicrobial peptides by other researchers. However, our previous work has shown that certain cationic antimicrobial peptides can have activity against this organism [27-29, 52, 53]. Experiments to introduce such antimicrobial peptides in the lung by Flick-Smith et al only modestly increased the time-to-death of mice infected with *F. tularensis* LVS [20]. In this work, we sought to develop another antimicrobial peptide that would be effective against *Francisella in vitro* and which would display *in vivo* activity against this infection.

Previously, our group identified C-terminal fragments of apolipoprotein C-1 from *Alligator mississippiensis*. These helical fragments, called Apo5 and Apo6, were found to have broad-spectrum activity against a variety of pathogens, including *Pseudomonas aeruginosa, Staphylococcus aureus*, and *Acinetobacter baumannii* [16, 19]. In general, these peptides had strong anti-*Francisella* activity in low-salt buffer, with several peptides exhibiting EC_50_ values under 3 µg/ml. When we tested these peptides against *F. tularensis* subspecies, it was found that Apo5 and Apo6 were generally less effective against these subspecies than against other Gram negative bacteria tested, with EC_50_ values ranging from low (∼6 µg/ml) against *F. tularensis* LVS to much higher (∼16 µg/ml) against *F. tularensis* NIH B38.

We designed the GATR series of peptides with the aim of improving upon the promising antimicrobial performance demonstrated by these peptides. Because there was no discernable difference in activity between Apo5 and Apo6 against the *F. tularensis* subspecies, we chose to focus on derivatives of the shorter Apo6. Little is known regarding the antimicrobial mechanisms and the associated specific interactions employed by these peptides, hence the GATR peptide variants (GATR-1 to −7) were designed based on incremental minor changes to the peptide sequences. Individually, these modifications were anticipated to minimally impact the peptide structural properties and preserve amino acid side-chain groups present in the parent peptide that may participate in critical interactions with bacterial targets such as the membrane or LPS. By substituting amino acids to increase peptide hydrophobicity and overall positive charge, we were able to create peptides with stronger *in vitro* and *in vivo* activity. All of the peptides, except GATR-4, exhibited superior performance over the parent peptide Apo6 against *F. tularensis* LVS under EC_50_ conditions. All the GATR variants, except GATR-1 and GATR-2, also demonstrated superior performance against *F. tularensis* NIH B38, the *F. tularensis* type strain, compared to parent peptide Apo6. The more substituted peptides, GATR-5, −6, and −7, began to show activity in cation-adjusted Mueller Hinton Broth, though a MIC could not be determined based upon the concentrations tested. The most efficacious peptides were also found to have stronger activity against *F. tularensis* SchuS4 compared to Apo6. Most notably, GATR-7 had a determinable MIC against *F. tularensis* SchuS4 at concentrations tested.

Our previous studies showed that the Apo6 peptide affected bacteria by disrupting the bacterial membrane, primarily through depolarization [19]. To examine if this was also the case with the synthetic peptides with *F. tularensis*, we examined membrane binding and disruption. DiSC_3_(5) measures depolarization and transient holes in a previously hyperpolarized membrane. It was found that as hydrophobicity and cationicity increase, so does depolarization activity. However, this is not the case when the ethidium bromide uptake assay was performed, which measures larger pores or disruption that allow the passage of ethidium bromide into the cell. While Apo6 shows significant membrane disruption at both 10 and 1 µg/ml tested, none of the other peptides show significant depolarization at 1 µg/ml. In general, the GATR peptides have a similar ΔRFU to Apo6 at 10 µg/ml. It is not clear why this occurs based on physico-chemical properties. The charge and hydrophobicity of Apo6 is generally much lower than that of the GATR peptides, but considering the greater antimicrobial efficacy of the GATR peptides, it appears that the pore-forming activity is less important to its antibacterial mechanism than the depolarization activity, which may suggest an intracellular target.

Some antimicrobial peptides, such as LL-37, bind to bacterial LPS [14]. In addition, some apolipoproteins have been shown to bind LPS [54, 55]. Our prior experiments had shown that Apo6 did not bind *E. coli* LPS (data not shown), and we found, similarly, that Apo6 does not significantly bind *F*. *tularensis* LVS LPS. The synthetic GATR peptides, however, were found to bind greater amounts of *F. tularensis* LVS LPS as hydrophobicity and cationicity increased, leveling off with GATR-6 and GATR-7. It is unclear if increasing LPS binding leads to increased depolarization, but it is possible that increased attraction between peptide and LPS allows higher-binding peptides to better associate with the membrane. It seems that there is no correlation, either positive or negative, between LPS binding and pore formation in the membrane.

After dropping one peptide (GATR-5) for toxicity issues, the 3 best performing peptides, GATR-3, GATR-6, and GATR-7, were tested for activity *in vivo*. Though GATR-7 had the best performance in MIC assays, initial studies in *G. mellonella* waxworms did not clearly show this peptide to be the best performer. Instead, GATR-3 and the parent peptide Apo6 saved more larvae from a lethal *F. tularensis* infection, though GATR-6 and GATR-7 also significantly rescued the waxworms. Because the *in vitro* and *in vivo* data together did not point to a clear front-runner peptide, we tested the three top-performing peptides (GATR-3, GATR-6, GATR-7) in a murine model infected with pulmonary tularemia. In initial studies, we tested Apo6 *in vivo* because of its strong performance in *G. mellonella* studies. However, this peptide was not effective in murine studies (data not shown). In a set of further experiments, we tested GATR-3, GATR-6, and GATR-7, as well as LL-37 and its D-enantiomer D-LL-37. We have previously shown that LL-37 is a highly effective peptide against *Francisella in vitro* [27, 29, 34-36]. LL-37 peptide has been previously tested in a murine pulmonary tularemia model [20]. Flick-Smith et al previously reported that when LVS-infected mice are treated with LL-37 via the intranasal route, the peptide significantly extended mean time to death, but did not ultimately rescue any mice. In our experiments, we treated at a higher concentration than Flick-Smith et al and also by a different route (via intraperitoneal injection), and similarly found that LL-37 had no effect on survival of infected mice. We also tested D-LL-37 *in vivo* because we have shown in *in vitro* that this chiral enantiomer is equally or more effective than the native peptide [35, 36]. In addition, it has the advantage of protease resistance, which should allow it to circulate in the body longer. When D-LL-37 was tested in this model, this peptide was also ineffective at rescuing infected mice or even prolonging mean time till death. Thus, LL-37 is not effective against a pulmonary-based tularemia infection when given systemically, in agreement with previous reports [20].

When the GATR peptides were tested in this model with a prophylactic treatment, it was found that both GATR-3 and GATR-6 peptide treatments significantly rescued mice infected with *F. tularensis* LVS. GATR-7 did not, though this peptide had the strongest activity in MIC assays, which are considered the gold standard for activity [19, 24]. In a second set of experiments, GATR-3 and GATR-6 were tested in larger groups without the prophylactic treatment. GATR-3 maintained its efficacy without the prophylactic treatment, while GATR-6 did not. This indicates that the pre-infection administration at Day −1 may have been important for the activity of GATR-6, which makes this peptide less favorable as a candidate to take forward for further testing.

Other groups have used peptides to treat a variety of bacterial infections in animal models with varying levels of success. Silva et al treated mice infected with *E. coli* and *S. aureus* with peptides derived the marine tunicate *Styela clava* and found that a single dose of 10 mg/kg yielded survival rates of 80-90% [56]. In another study, it was found that a single 80-200 mg/kg dose boosted survival of rainbow trout infected with *Yersinia ruckeri* from 20% to 70% [57]. Additionally, mice infected with *Bacillus anthracis* spores were treated with a single dose of 1 mg/kg synthetic protease-resistant peptides, and survival was boosted to 20-30% [58]. Thus, 3-4 doses of 5 mg/kg GATR-3 yielding 50-60% survival in infected mice compares favorably with the results of other trials. The dosage of GATR-3 in this study is also comparable to levofloxacin (5 mg/kg vs. 3 mg/kg), though GATR-3 did not rescue all mice in the treated cohort. Larger doses of GATR-3 may increase efficacy of the peptide; however, we must first study safety of these larger doses in mice. Based on the treatment used in the study, a preliminary dosage for human infection can be inferred using guidelines put forth by the FDA [59]. The dosage of 5 mg/kg in mice would convert to approximately 0.4 mg/kg in humans with a dose of 24 mg presuming a 60 kg human.

*F. tularensis* disseminates from the lung to the liver and spleen during infection [20]. GATR-3 was tested in an organ burden study to examine whether bacterial burden reduction was the cause for *in vivo* activity. Although *Francisella* burden was reduced in the lungs of GATR-3-treated mice, it was not significantly reduced in the liver or spleen. Further studies are needed to determine whether GATR-3 clears the infection directly as its mode of action *in vivo* or whether it activates the host immune response to promote survival. Due to its strong and consistent performance, GATR-3 has strong potential as an anti-tularemia peptide and will be the subject of further pre-clinical development to determine its pharmacokinetic and pharmacodynamics properties.

## Acknowledgements

SMB, AK, BMB, and MVH were supported by HDTRA1-12-C-0039 “Translational Peptide Research for Personnel Protection.” We would like to acknowledge Evelyn Hrifko, Kajal Gupta, and Brittany Heath for technical assistance. We would also like to thank Joel Schnur for helpful discussions.

## References

1. Gürcan, Ş., Epidemiology of Tularemia. Balkan medical journal, 2014. 31(1): p. 3–10.

2. Propst, C.N., et al., Francisella philomiragia Infection and Lethality in Mammalian Tissue Culture Cell Models, Galleria mellonella, and BALB/c Mice. Front Microbiol, 2016. 7: p. 696.

3. Maurin, M., Francisella tularensis as a potential agent of bioterrorism? Expert Rev Anti Infect Ther, 2015. 13(2): p. 141–4.

4. Notifiable Diseases and Mortality Tables. Morbidity and Mortality Weekly Report, 2018. 66(52).

5. Control, E.C.f.D.P.a., Annual Epidemiological Report 2016 – Tularaemia. 2016, ECDC: Stockholm.

6. Stevens, D.L., et al., Practice guidelines for the diagnosis and management of skin and soft tissue infections: 2014 update by the Infectious Diseases Society of America. Clin Infect Dis, 2014. 59(2): p. e10–52.

7. Li, Y., et al., LPS remodeling is an evolved survival strategy for bacteria. Proc Natl Acad Sci U S A, 2012. 109(22): p. 8716–21.

8. Llewellyn, A.C., et al., NaxD is a deacetylase required for lipid A modification and Francisella pathogenesis. Mol Microbiol, 2012. 86(3): p. 611–27.

9. Brikman, D.I. and O.D. Zakhlebnaia, [Sensitivity of various races of the tularemia microbe to erythromycin, oleandomycin and polymyxin]. Antibiotiki, 1976. 21(2): p. 134–6.

10. Pavlovich, N.V., et al., [Detection of persistent resistance to antibacterial drugs in various strains of Francisella tularensis]. Antibiot Khimioter, 1992. 37(10): p. 29–31.

11. Nakatsuji, T. and R.L. Gallo, Antimicrobial Peptides: Old Molecules with New Ideas. J Invest Dermatol, 2012. 132(3): p. 887–895.

12. Pane, K., et al., A new cryptic cationic antimicrobial peptide from human apolipoprotein E with antibacterial activity and immunomodulatory effects on human cells. FEBS J, 2016. 283(11): p. 2115–31.

13. Pane, K., et al., Antimicrobial potency of cationic antimicrobial peptides can be predicted from their amino acid composition: Application to the detection of “cryptic” antimicrobial peptides. J Theor Biol, 2017. 419: p. 254–265.

14. Turner, J., et al., Activities of LL-37, a cathelin-associated antimicrobial peptide of human neutrophils. Antimicrob Agents Chemother, 1998. 42(9): p. 2206–14.

15. Dürr, U.H.N., U.S. Sudheendra, and A. Ramamoorthy, LL-37, the only human member of the cathelicidin family of antimicrobial peptides. Biochimica et Biophysica Acta (BBA) - Biomembranes, 2006. 1758(9): p. 1408–1425.

16. Bishop, B.M., et al., Bioprospecting the American Alligator (Alligator mississippiensis) Host Defense Peptidome. PLoS One, 2015. 10(2): p. e0117394.

17. Juba, M.L., et al., Large Scale Discovery and De Novo-Assisted Sequencing of Cationic Antimicrobial Peptides (CAMPs) by Microparticle Capture and Electron-Transfer Dissociation (ETD) Mass Spectrometry. J Proteome Res, 2015. 14(10): p. 4282–95.

18. Bishop, B.M., et al., Discovery of Novel Antimicrobial Peptides from Varanus komodoensis (Komodo Dragon) by Large-Scale Analyses and De-Novo-Assisted Sequencing Using Electron-Transfer Dissociation Mass Spectrometry. J Proteome Res, 2017.

19. Barksdale, S.M., et al., Peptides from American alligator plasma are antimicrobial against multi-drug resistant bacterial pathogens including Acinetobacter baumannii. BMC Microbiol, 2016. 16(1): p. 189.

20. Flick-Smith, H.C., et al., Assessment of antimicrobial peptide LL-37 as a post-exposure therapy to protect against respiratory tularemia in mice. Peptides, 2013. 43: p. 96–101.

21. Gautier, R., et al., HELIQUEST: a web server to screen sequences with specific alpha-helical properties. Bioinformatics, 2008. 24(18): p. 2101–2.

22. Wang, G., X. Li, and Z. Wang, APD3: the antimicrobial peptide database as a tool for research and education. Nucleic Acids Res, 2016. 44(D1): p. D1087–93.

23. Georgi, E., et al., Standardized broth microdilution antimicrobial susceptibility testing of Francisella tularensis subsp. holarctica strains from Europe and rare Francisella species. J Antimicrob Chemother, 2012. 67(10): p. 2429–33.

24. Institute, C.a.L.S., Performance Standards for Antimicrobial Susceptibility Testing, in M100-S22. 2012, CLSI: Wayne, PA, USA.

25. Giacometti, A., et al., In vitro susceptibility tests for cationic peptides: comparison of broth microdilution methods for bacteria that grow aerobically. Antimicrob Agents Chemother, 2000. 44(6): p. 1694–6.

26. Han, S., B.M. Bishop, and M.L. van Hoek, Antimicrobial activity of human beta-defensins and induction by Francisella. Biochem Biophys Res Commun, 2008. 371(4): p. 670–4.

27. Amer, L.S., B.M. Bishop, and M.L. van Hoek, Antimicrobial and antibiofilm activity of cathelicidins and short, synthetic peptides against Francisella. Biochem Biophys Res Commun, 2010. 396(2): p. 246–51.

28. Kaushal, A., et al., Antimicrobial activity of mosquito cecropin peptides against Francisella. Dev Comp Immunol, 2016. 63: p. 171–180.

29. Chung, M.C., S.N. Dean, and M.L. van Hoek, Acyl carrier protein is a bacterial cytoplasmic target of cationic antimicrobial peptide LL-37. Biochem J, 2015. 470(2): p. 243–53.

30. Bland, J.M., et al., All-D-cecropin B: synthesis, conformation, lipopolysaccharide binding, and antibacterial activity. Mol Cell Biochem, 2001. 218(1-2): p. 105–11.

31. Barksdale, S.M., E.J. Hrifko, and M.L. van Hoek, Cathelicidin antimicrobial peptide from Alligator mississippiensis has antibacterial activity against multi-drug resistant Acinetobacter baumanii and Klebsiella pneumoniae. Dev Comp Immunol, 2017.

32. Sprynski, N., E. Valade, and F. Neulat-Ripoll, Galleria mellonella as an infection model for select agents. Methods Mol Biol, 2014. 1197: p. 3–9.

33. Dean, S.N. and M.L. van Hoek, Screen of FDA-approved drug library identifies maprotiline, an antibiofilm and antivirulence compound with QseC sensor-kinase dependent activity in Francisella novicida. Virulence, 2015. 6(5): p. 487–503.

34. M.C. Chung, E., et al., Komodo dragon-inspired synthetic peptide DRGN-1 promotes wound-healing of a mixed-biofilm infected wound. npj Biofilms and Microbiomes, 2017. 3(1): p. 9.

35. Dean, S.N., B.M. Bishop, and M.L. van Hoek, Natural and synthetic cathelicidin peptides with anti-microbial and anti-biofilm activity against Staphylococcus aureus. BMC Microbiol, 2011. 11: p. 114.

36. Dean, S.N., B.M. Bishop, and M.L. van Hoek, Susceptibility of Pseudomonas aeruginosa Biofilm to Alpha-Helical Peptides: D-enantiomer of LL-37. Front Microbiol, 2011. 2: p. 128.

37. Travis, S.M., et al., Bactericidal Activity of Mammalian Cathelicidin-Derived Peptides. Infection and Immunity, 2000. 68(5): p. 2748–2755.

38. Epand, R.F., et al., Depolarization, bacterial membrane composition, and the antimicrobial action of ceragenins. Antimicrob Agents Chemother, 2010. 54(9): p. 3708–13.

39. Brandenburg, K., et al., Peptides with dual mode of action: Killing bacteria and preventing endotoxin-induced sepsis. Biochim Biophys Acta, 2016. 1858(5): p. 971–9.

40. Travkova, O.G., H. Moehwald, and G. Brezesinski, The interaction of antimicrobial peptides with membranes. Adv Colloid Interface Sci, 2017.

41. Sun, Y. and D. Shang, Inhibitory Effects of Antimicrobial Peptides on Lipopolysaccharide-Induced Inflammation. Mediators Inflamm, 2015. 2015: p. 167572.

42. Coorens, M., et al., Interspecies cathelicidin comparison reveals divergence in antimicrobial activity, TLR modulation, chemokine induction and regulation of phagocytosis. Sci Rep, 2017. 7: p. 40874.

43. Nagaoka, I., et al., Antibacterial cathelicidin peptide CAP11 inhibits the lipopolysaccharide (LPS)-induced suppression of neutrophil apoptosis by blocking the binding of LPS to target cells. Inflamm Res, 2004. 53(11): p. 609–22.

44. Tack, B.F., et al., SMAP-29 has two LPS-binding sites and a central hinge. Eur J Biochem, 2002. 269(4): p. 1181–9.

45. Vinogradov, E., M.B. Perry, and J.W. Conlan, Structural analysis of Francisella tularensis lipopolysaccharide. Eur J Biochem, 2002. 269(24): p. 6112–8.

46. Gunn, J.S. and R.K. Ernst, The structure and function of Francisella lipopolysaccharide. Ann N Y Acad Sci, 2007. 1105: p. 202–18.

47. Jones, C.L., et al., Subversion of host recognition and defense systems by Francisella spp. Microbiol Mol Biol Rev, 2012. 76(2): p. 383–404.

48. Aperis, G., et al., Galleria mellonella as a model host to study infection by the Francisella tularensis live vaccine strain. Microbes Infect, 2007. 9(6): p. 729–34.

49. Blower, R.J., S.G. Popov, and M.L. van Hoek, Cathelicidin peptide rescues G. mellonella infected with B. anthracis. Virulence, 2017: p. 1–7.

50. Petersen, J.M., et al., Direct isolation of Francisella spp. from environmental samples. Lett Appl Microbiol, 2009. 48(6): p. 663–7.

51. Kanistanon, D., et al., Role of Francisella lipid A phosphate modification in virulence and long-term protective immune responses. Infect Immun, 2012. 80(3): p. 943–51.

52. van Hoek, M.L., Biofilms: an advancement in our understanding of Francisella species. Virulence, 2013. 4(8): p. 833–46.

53. Chung, M.C., et al., Chitinases are negative regulators of Francisella novicida biofilms. PLoS One, 2014. 9(3): p. e93119.

54. Emancipator, K., G. Csako, and R.J. Elin, In vitro inactivation of bacterial endotoxin by human lipoproteins and apolipoproteins. Infect Immun, 1992. 60(2): p. 596–601.

55. Beck, W.H.J., et al., Apolipoprotein A–I binding to anionic vesicles and lipopolysaccharides: Role for lysine residues in antimicrobial properties. Biochimica et Biophysica Acta (BBA) - Biomembranes, 2013. 1828(6): p. 1503–1510.

56. Silva, O.N., et al., An anti-infective synthetic peptide with dual antimicrobial and immunomodulatory activities. Sci Rep, 2016. 6: p. 35465.

57. Chettri, J.K., et al., Antimicrobial peptide CAP18 and its effect on Yersinia ruckeri infections in rainbow trout Oncorhynchus mykiss (Walbaum): comparing administration by injection and oral routes. J Fish Dis, 2016.

58. Teyssieres, E., et al., Proteolytically Stable Foldamer Mimics of Host-Defense Peptides with Protective Activities in a Murine Model of Bacterial Infection. J Med Chem, 2016. 59(18): p. 8221–32.

59. Guidance for Industry: Estimating the Maximum Safe Starting Dose in Initial Clinical Trials for Therapeutics in Adult Healthy Volunteers F.a.D. Administration, Editor. 2005: Rockville, MD.

